# Drug binding sites in the multidrug transporter MdfA in detergent solution and in lipid nanodiscs

**DOI:** 10.1101/2020.08.31.275321

**Authors:** T. Bahrenberg, E. H. Yardeni, A. Feintuch, E. Bibi, D. Goldfarb

## Abstract

MdfA, a member of the major facilitator superfamily (MFS), is a multidrug/proton antiporter from *E. coli* that has been considered a model for secondary multidrug (Mdr) transporters. Its transport mechanism, driven by a proton gradient, is associated with conformational changes, which accompany the recruitment of drugs and their release. In this work, we applied double-electron electron resonance (DEER) spectroscopy to locate the binding site of one of its substrates, tetraphenylphosphonium (TPP) within available crystal structures. We carried out Gd(III)-nitroxide distance measurements between MdfA labeled with a Gd(III) tag and the TPP analog mito-TEMPO (bearing the nitroxide moiety). Data were obtained both for MdfA solubilized in detergent micelles (n-dodecyl-β-D-maltopyranoside (DDM)), and reconstituted into lipid nanodiscs (ND). For both DDM and ND, the average position of the substrate at a neutral pH was found to be close to the ligand position in the I_f_ (inward facing) crystal structure, with the DDM environment exhibiting a somewhat better agreement than the ND environment. We therefore conclude that the I_f_ structure provides a good description for substrate-bound MdfA in DDM solution, while in ND the structure is slightly modified. A second binding site was found for the ND sample situated at the cytoplasmic side, towards the end of transmembrane helix 7 (TM7). In addition, we used DEER distance measurements on Gd(III) doubly labeled MdfA to track conformational changes within the periplasmic and cytoplasmic sides associated with substrate binding. We detected significant differences in the periplasmic side of MdfA, with the ND featuring a more closed conformation than in DDM, in agreement with earlier reports. The addition of TPP led to a noticeable conformational change in the periplasmic face in ND, attributed to a movement of TM10. This change was not observed in DDM.

**Statement of Significance:** MdfA is multidrug transporter from *E. coli*, which exhibits multidrug efflux activities with an unusually broad spectrum of drug specificities. While it has been established that solute transport by similar transporters is coupled to significant conformational changes, previous studies raised the possibility that this is not the case for MdfA. Moreover, it is not clear how MdfA functionally accommodates chemically dissimilar substrates. Towards resolving these open questions, we used double-electron electron resonance distance measurements to determine the binding site of a spin labeled drug analog within available crystal structures of MdfA and to examine how MdfA responds conformationally to drug binding. Moreover, we explored how these two are affected by the media, detergent micelles vs lipid nanodiscs.

## Introduction

MdfA, a member of the major facilitator superfamily (MFS), is a multidrug/proton antiporter from *E. coli* that has been considered a model for secondary multidrug (Mdr) transporters.(1) MdfA is capable of transporting a variety of structurally dissimilar compounds out of the cell, including antibiotics like chloramphenicol.(2) This transport is driven by a proton gradient, and in each transport cycle MdfA imports one proton into the cell.(2) Similar to most of MFS antiporters, MdfA folds in the canonical, basic MFS fold which consists of 12 transmembrane (TM) helices.(3, 4) Previous studies suggest that MdfA is unusually flexible (1), a fact that makes crystallization difficult. The first high-resolution X-ray structure of MdfA was published only in 2015 in an inward facing conformation (I_f_, PDB 4ZP0),(5) and more recently, another crystal structure that depicts MdfA in an outward open conformation (O_o_, PDB 6GV1) has been published.(6) Both crystal structures, however, do not represent MdfA in its native state. The I_f_ crystal structure was determined from the non-functional mutant Q131R with bound deoxycholate and other ligands(5), while for the more recent O_o_ structure, crystallization was facilitated by a Fab antibody fragment attached to the cytoplasmic side of the protein.(6)

The solution structure of MdfA solubilized in detergent at neutral pH, was explored by double electron-electron resonance (DEER) measurements on a series of double cysteine mutants of MdfA labeled with two different spin labels – a pair of nitroxide labels (NO) or a pair of C2-Gd labels (see Fig. 1a).(7) Both labels, which are dissimilar in their structures, sizes, charge and hydrophobicities, revealed the flexibility of detergent-solubilized MdfA, and showed that the periplasmic side has a similar structure to the O_o_ conformation reported by the 6GV1 crystal structure, though somewhat more closed, and is quite different from the I_f_ crystal structure. For the cytoplasmic face, the results show only a slight similarity to the O_o_ with a more open conformation.

**Figure 1.**
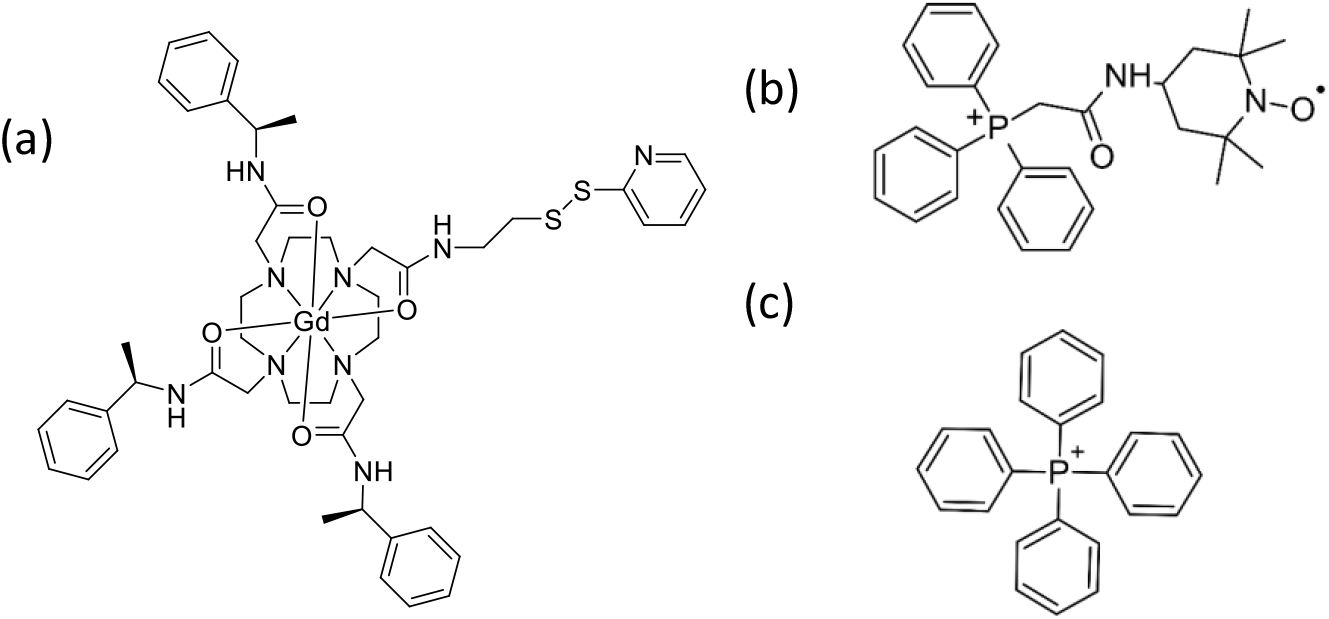
Chemical structure of (a) Gd-C2 (b) mito-TEMPO, (c) TPP.

A model of the transport process suggested that MdfA relies on the alternate-access mechanism, in which the protein is open towards the cytoplasm, undergoes a substrate-induced structural rearrangement that closes the cytosolic region and opens the protein towards the periplasm, and vice versa.(8) Relating these steps to published crystal structures suggests that the O_o_ conformation presumably captures the state in which substrates are released in the periplasm. On the other hand, the ligand occluded I_f_ conformation is assumed following deprotonation. MdfA’s transport capabilities include neutral, zwitterionic and cationic cytotoxic compounds such as ethidium bromide, tetraphenylphosphonium (TPP) bromide, ciprofloxacin, and chloramphenicol.(9) Drug/proton exchange requires a site that can be protonated, which in MdfA has been identified as Asp34 that together with Glu26 are essential for transporting cationic drugs.(10) Results from biochemical studies suggested that a substrate must be released prior to binding of a proton, so there is a competition between binding a substrate and binding a proton.(11) Mechanistically, in the I_f_ conformation residue Asp34 gets deprotonated, enabling the binding of a substrate in the multidrug recognition pocket. A conformational change towards the O_o_ conformation is triggered by this binding. The substrate is released to the periplasm and, consequently, the Asp34 residue becomes exposed. Following protonation, the protein undergoes a second conformational change towards the I_f_ structure and the cycle is complete. Interestingly, this mechanism implies that the antiporter cannot export cationic substrates at an external alkaline pH above 7.6.(1)

Recently, MdfA has been studied in phospholipid bilayers, more specifically in nanodiscs (ND) and native membrane by DEER spectroscopy and biochemical cysteine cross-linking.(12) The study reported that in this environment, MdfA assumes a relatively flexible, outward-closed/inward-closed (O_c_/I_c_) conformation,(12) which is different from the observation in detergent.(7) Moreover, the data show that neither the substrate TPP nor protonation induces large scale conformational changes as expected from the above description. In turn, the authors identified a substrate-responsive lateral gate, which is open toward the inner leaflet of the membrane and closes upon drug binding. This recent work suggested a modified model for the functional conformational cycle of MdfA that does not invoke canonical structural elements of alternating access. So far, the experimental structural studies of MdfA in detergent or in membrane mimics, focused on the conformational changes of MdfA upon substrate binding(7, 8, 12, 13) but not directly on the location of the substrate within the structure.

In the present work, we focus on the location of TPP within the structure of MdfA. We investigated both MdfA solubilized in DDM (n-Dodecyl-B-D-Maltoside) detergent and reconstituted in lipid ND, looking for indications that structural differences may be induced by the lipid environment. We labeled MdfA with a Gd(III) spin label, C2-Gd (Fig. 1a), as used earlier(7) and added to it an NO labeled TPP analogue, mito-TEMPO (see Fig. 1b,c). We carried out W-band (95 GHz) Gd(III)-NO DEER measurements on three singly labeled C2-Gd sites on the periplasmic side in the presence of bound mito-TEMPO and obtained the Gd(III)-NO distance distributions. The advantage of using Gd(III)-NO DEER, as opposed to NO-NO or Gd(III)-Gd(III) DEER, is the ability to track protein-substrate distances for weak binding with minimal interferences from overlapping EPR signals from unbound substrate. Using DEER derived distance distributions and triangulation, we located the NO group of mito-TEMPO within the I_f_ structure, which was solved with several ligands and a substrate(5), and the O_o_ structure(6). The location of the substrate agreed rather well with the I_f_ and not the expected O_o_ structure, and the DDM results showed better agreement than the ND results. This suggests that the I_f_ structure, which was obtained from MdfA solubilized in DDM, is a good representation for substrate bound MdfA solubilized in DDM and less so for ND. In addition, using Gd(III)-Gd(III) DEER on doubly labeled MdfA to track conformational changes upon TPP binding, we identified significant conformational changes on the periplasmic side of MdfA in ND but not in DDM.

## Experimental

Gd-C2 was obtained from Bim Graham and synthesized as described earlier(14). Mito-TEMPO chloride was purchase from Sigma-Aldrich and used without further purification. TPP bromide was purchased from Fluka. Preparation of the double cysteine-less (CL)-MdfA mutants A128C/S280C and L168C/S310C, the single CL mutants L101C, S310C and L168C, their solubilization in 1.1% n-dodecyl-β-D-maltopyranoside (DDM) detergent and their labeling with Gd-C2 was done as described earlier.(7) The MdfA activity of all single and double cysteine mutants was confirmed earlier.(7)

### Preparation of MdfA reconstituted in ND

#### The MSP protein

Here we followed the protocol described in (12, 15). *E. coli* BL21(DE3) transformed with plasmid pET-28a(+) encoding MSP1D1E3 were grown in Terrific Broth (TB) supplemented with 30 µg/ml Kanamycin at 37 °C to A_600_ of 2.5-3, and induced using 1 mM IPTG. After 4 h of expression, cells were harvested and resuspended in lysis buffer (50 mM sodium phosphate buffer, 1 % Triton X-100), supplemented with 1 tablet of Complete Mini, EDTA-free Protease Inhibitor Cocktail (Roche) and 1 mM PMSF. Cells were then lysed by sonication (two cycles, 8 min at 30% amplitude, 15 s on, 15 s off, using Sonics Ultra Cell). The suspension was ultracentrifuged at 30,000 g for 30 min to remove non-lysed material. The protein was then purified by Ni-NTA affinity chromatography and eluted with 300 mM Imidazole in a tris-buffer (40 mM Tris, 300 mM NaCl, 300 mM Imidazole, pH 8.0). Finally, the protein was desalted using a HP Desalting column (GE Healthcare) into 50 mM Tris/MES buffer (pH 7.5), concentrated to a volume of around 2 ml (∼15 mg/ml) and stored at 4 °C.

#### Phospholipids

Phospholipids (*E. coli* polar lipid extract, Avanti Polar Lipids) were prepared by dissolving the powder in CHCl_3_/MeOH (3:2) and subsequent removal of the solvent in vacuo. Lipids were then hydrated by addition of 20 mM Tris-HCl, pH 7.5, 0.12 M NaCl (15mg/ml) and subjected to ten cycles of freezing and thawing in liquid nitrogen, followed by sonication. Lipids were then flash frozen and stored at −80°C.

The desired MdfA single or double mutant was grown, purified, and labeled as described earlier.(7) Following labeling, the sample was loaded on a Superdex 200 10/30 GL (GE) gel filtration column, eluted, and appropriate fractions combined after verification by SDS-PAGE. The protein volume was reduced to 0.5 ml using a Vivaspin concentrator with a cutoff of 10 kDa and fresh MSP (MdfA:MSP 1:10), lipids (MSP:Lipids 1:60, *E. coli* polar lipid extract) and 0.1 M Sodium Cholate (Lipids:Detergent 1:5, where detergent refers to the total detergent concentration in the final sample, including DDM) were added. After 1 h of incubation at 4 °C, Biobeads (Bio-rad) were added stepwise: 100 mg per 1 ml of sample after 1 h, 100 mg after 2 h, 200 mg after 3 h and another 200 mg after overnight incubation. After removal of the beads by centrifugation the sample was concentrated using a 100 kDa Vivaspin concentrator, loaded on a Superdex 200 10/30 GL gel filtration column and eluted using 20 mM Tris-HCl, pH 7.5, 120 mM NaCl, 10% glycerol. Following elution, fractions containing MdfA, embedded in nanodiscs were combined, as determined by SDS-PAGE for the presence of MSP and MdfA (see Fig. S1).

### Preparation of MdfA + TPP or mito-TEMPO samples for DEER

TPP bromide was added in excess (1 mM) to doubly labeled MdfA, mixed and the mixture incubated for 15 minutes on ice. Samples containing mito-TEMPO were prepared by adding 1, 2, or 5 equivalents of mito-TEMPO from a 1 mM stock solution in 20 mM Tris-HCl buffer (pH 7.5) to the protein sample, followed by mixing and 20 min incubation on ice.

All EPR samples were in 20 mM Tris-HCl, pH 7.5, 120 mM NaCl buffer D_2_O glycerol-d_8_, (9:1 v/v). Protein concentration was determined by measuring the absorbance at 280 nm, *A*_280_ (1 mg/mL ∼ 2.1 *A*_280_) and the protein was concentrated to a final concentration of 20 – 50 μM using a 100K MWCO concentrator (Amicon). Finally the sample was loaded in 0.8 o.d. 0.6 i.d mm quartz capillaries, flash frozen in liquid nitrogen and kept until measured.

### Mito-TEMPO binding constant measurements

Binding of mito-TEMPO was performed as described earlier(16) with small modifications. Purified protein (2 µg/sample) was mixed with Ni-TNA (talon) resin (10 µl of suspended resin/sample) in buffer A (20 mM Tris-HCl, pH 8, 0.5 M NaCl, 10% glycerol, 0.1% DDM, 5 mM imidazole), and incubated at 4°C for 1 h, with gentle agitation. Unbound protein was then discarded by centrifugation (2 min, 700 g) and the resin was resuspended in 100 μl/sample of buffer (20 mM Tris-HCl, pH 8.0, 0.5 M NaCl, 0.1% DDM). Samples were then mixed with 100 μl/sample of buffer C, supplemented with 2 µM of [^3^H]TPP (two-fold concentration) and mito- TEMPO at concentrations ranging from 0-80 µM (two fold concentration). The final mixture, containing 1 µM of [^3^H]TPP (1 Ci/mmol) and 0-40 µM of mito-TEMPO, was then incubated at 4°C for 15 min with continuous tilting. The resin (180 μl) was then transferred to a Promega Wizard minicolumn, placed on a microcentrifuge tube, and centrifuged at >10,000 g for 20 sec. Unbound material (flow-through) was discarded, and the resin was resuspended in 100 μl of buffer D (20 mM Tris-HCl, pH 7.0, 0.5 M NaCl, 0.1% DDM, 350 mM imidazole) and removed to scintillation vials. Scintillation fluid (5 ml) was added to the samples and then measured in a beta-counter for 1 min. To determine the inhibition constant of mito- TEMPO (K_i_), the data was fitted to the equation B=B_max_-B_max_([I]/[I]+K_i_) using nonlinear regression, where B equals to bound TPP (µM), B_max_ equals to the maximum of bound TPP (µM), and [I] refers to the concentration of mito-TEMPO (µM). Calculation of K_i_ and nonlinear regression was performed using MATLAB Curve Fitting toolbox.

### Spectroscopic measurements

Continuous-wave EPR experiments of the Gd(III)-NO samples were performed at X-band (9.4 GHz) on a Bruker E500 spectrometer at room temperature. Modulation amplitude was 1 G, Time constant 164 ms, conversion time 40.96 ms, and sweep time 41.94 ms. All pulsed EPR experiments were performed on a home-built 95 GHz (W-band) EPR spectrometer equipped with two microwave channels and an Arbitrary Waveform Generator (AWG). The pulse sequence used for the echo detected EPR (ED-EPR) was *π/2 – τ – π – τ – echo*. For optimized the observation the NO signal the conditions were as follows: 50 K: *t*_π/2,π_ = 20/40 ns, τ = 600 ns, shot repetition rate 5 ms. For optimizing Gd(III) signal : 10 K, *t*_π/2,π_ = 20/40 ns, τ = 600 ns, shot repetition rate 500 μs (see Fig. S2).

#### Gd(III)-Gd(III) DEER

DEER experiments were recorded at 10 K with the standard dead-time free DEER sequence (π/2_*v*obs_ – τ_1_ – π_*v*obs_ – (τ_1_+t) – π_*v*pump_ – (τ_2_-t) – π_*v*obs_ – τ_2_ – echo)(17) where the pump pulse was replaced with a linear chirp(18, 19), referred to as AWG DEER or rDEER(20), where the shaped pump pulse is swept between the primary echo and the first refocusing pulse. An 8-step phase cycling on the observer pulses was used; the observed pulse was set at 94.9 GHz, which corresponds to maximum echo intensity with pulse lengths: *t*_π/2,π_ = 15/30 ns. Two consecutive pump pulses set at 94.5-94.8 GHz and 95-95.3 GHz with pulse lengths of 96 ns each were applied. Most data were collected with rDEER where τ_2_=2.2 µs and the delay that determines the length of the DEER evolution time, τ_1_, varied from sample to sample. When AWG DEER was applied τ_1_= 1 µs was used and τ_2_ varied according to the DEER evolution time needed (or possible).

#### Gd(III)-NO DEER

Gd(III)-NO rDEER experiments were performed at 10 K by tuning the resonator at 94.9 GHz and setting the field maximum of the NO spectrum to 94.95 GHz. A single, 128 ns long linear chirp was used as a pump pulse from 94.9 – 95.1 GHz while the observe sequence was placed at 94.8 GHz. An 8-step phase cycling was used, shot repetition time was 200 μs. Here τ_2_ was in the range of 1.0-2.2, depending on the sample. Fig. S2 shows the DEER set up for the Gd(III)-Gd(III) and Gd(III)-NO DEER. The echo detection observe pulses were optimized for Gd(III) with lengths of 12.5 or 15 ns for the π/2 pulses and 25 or 30 ns for the π. The repetition rate was 0.2-0.5 ms, to ensure saturation of the NO signal.

#### Data analysis

DEER data were analyzed using Tikhonov regularization with the DeerAnalysis 2018 software package.(21) The background decay was fitted with a dimension of 3 and default values were used for the validation process including noise addition.

The calculated Gd(III)-Gd(III) distance distribution were carried out as described earlier(7) and the same program was used to calculate the average position of the Gd(III) in the Gd-C2 labels. These coordinates were used in the triangulation to determine the position of the NO radical in the mito-TEMPO in the I_f_ and O_o_ crystal structures.

## Results

### The effect of the environment and substrate on the conformation of MdfA

We first explored the effect of the environment, namely ND vs DDM micelles, on the conformation of MdfA. We chose two cysteine-less (CL) MdfA double mutants, A128C/S280C and L168C/S310C designated as TM4^128^ - TM8^280^ and TM5^168^ - TM10^310^, respectively. This nomenclature, used also in our earlier publication(7), highlights their location in terms of the MdfA TM helices. The first pair probes the cytoplasmic side of MdfA, whereas the second pair examines the periplasmic side (see Fig. 2).

**Figure 2.**
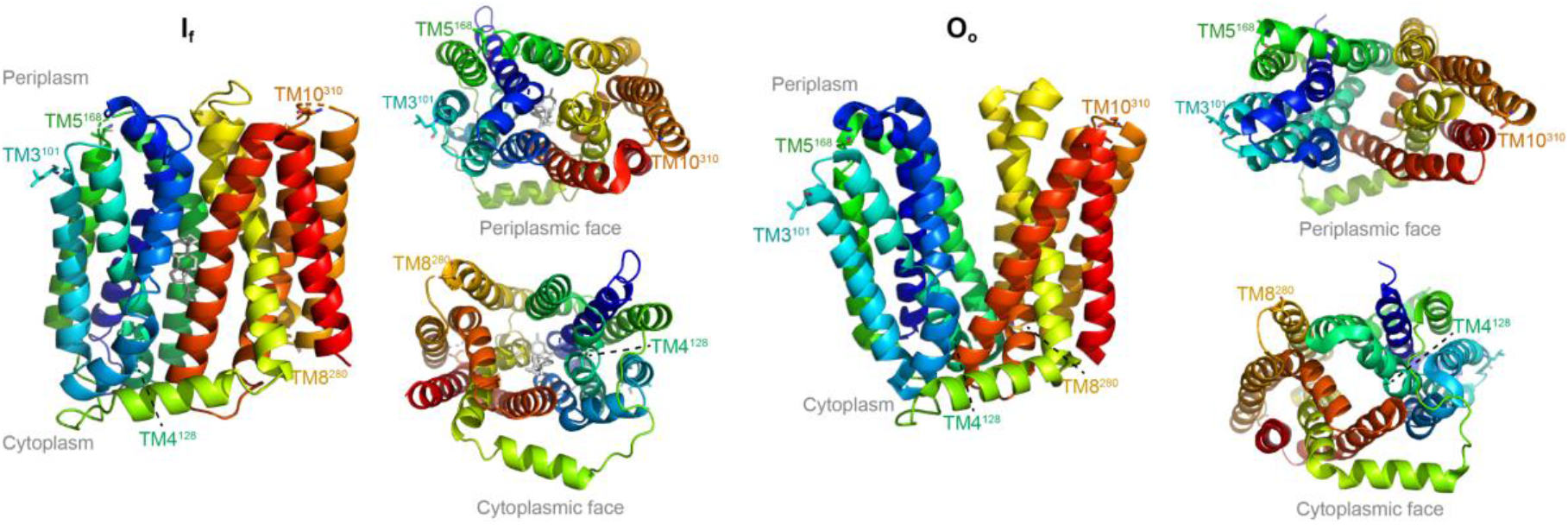
Labeling positions in MdfA, illustrated on several views, on the I_f_ (PDB 4ZP0(5)) and O_o_ (PDB 6GV1(6)) crystal structures. The substrate deoxycholate is shown as grey sticks in the I_f_ structure and the labeling residues are shown as sticks with colors matching the color of the TM helix they are part of. The periplsamic and cytoplasmic sides of the membrane are indicated.

DEER form factors and the corresponding distance distributions of TM4^128^ - TM8^280^ and TM5^168^ - TM10^310^ in DDM and ND, presented in Fig. 3, show that while the ND environment leads to a significant shortening of the distance between the periplasmic labels by ∼ 1 nm, as compared to DDM micelles, no change was observed for the cytoplasmic pair. Comparison of NO-NO distance measurements of the cytoplasmic pair TM1^20^-TM8^280^ in DDM and ND revealed a small change in the maxima of the distance distribution (0.37 nm)(7, 12) consistent with our observation. For the periplasmic pair TM3^101^-TM11^373^, the ND distance was shorter by 0.82 nm, whereas for the TM5^163^-TM10^307^, TM5^163^-TM11^373^ and TM5^168^-TM11^373^ pairs the shortening was considerably milder, 0.13-0.35 nm.(7, 12) We also compared the obtained distances to those predicted from the I_f_ and O_o_ crystal structures (PDB 4ZP0(5) and 6GV1(6)). For TM4^128^ - TM8^280^ the maxima of the predicted distance distribution is 2.8 nm for both structures(7), which is in a good agreement with the experimental maxima of 2.85 nm for the DDM sample and 2.9 nm for the ND sample. For TM5^168^ – TM10^310^ the predicted distance derived from the I_f_ structure (4.35 nm) agrees with the DDM experimental results, whereas the O_o_ predicts a maxima at 5.45 nm, which is in disagreement with both DDM and ND results.

**Figure 3.**
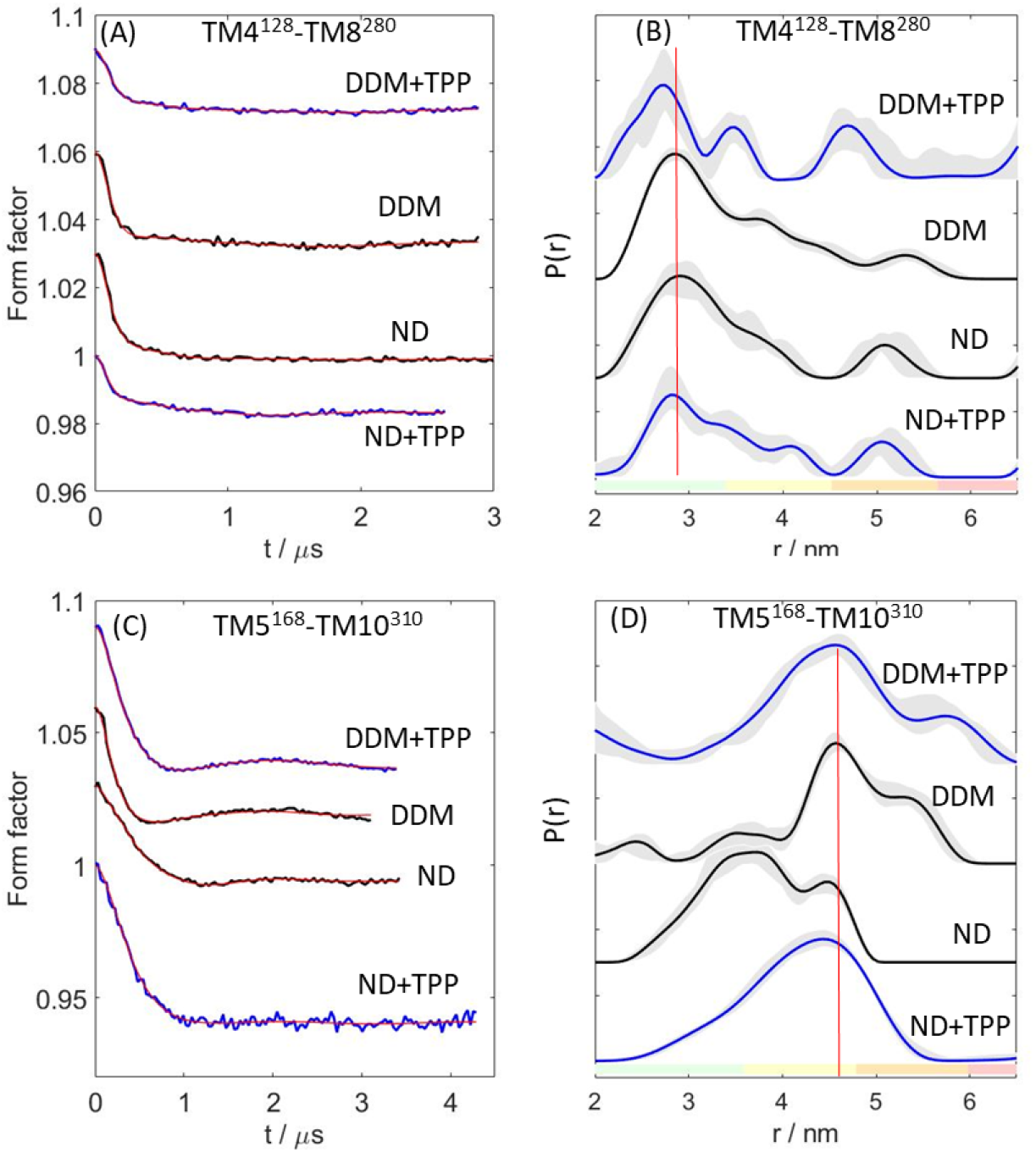
Gd(III)-Gd(III) DEER results on (A,B) TM4^128^ - TM8^280^ and (C,D) TM5^168^ – TM10^310^ in DDM and ND with and without the drug TPP. (A,C) DEER form factors obtained from the primary DEER after background correction and (B,D) the corresponding distance distribution with color coding of the reliability regions as defined in DeerAnalysis. (Pale green: the shape of the distance distribution is reliable. Pale yellow: the mean distance and distribution width are reliable. Pale orange: the mean distance is reliable. Pale red: long-range distance contributions may be detectable, but cannot be quantified.) The light grey regions, representing the uncertainty range of the data, show alternative distributions obtained by varying the parameters of the background correction as calculated by the validation tool in the DeerAnalysis software package. The vertical red lines are drawn to highlight the shift of the distance distribution. The primary DEER data are shown in Fig. S3

Next, we looked at the effect of TPP addition on the distance distributions of TM4^128^ - TM8^280^ and TM5^168^ – TM10^310^ in DDM and ND. For the cytoplasmic TM4^128^ - TM8^280^ pair, the addition of the drug did not lead to any significant change in either environments, whereas for the periplasmic TM5^168^ – TM10^310^, a significant shift was observed only for the ND sample. The maxima of the distance distribution shifted from 3.6 nm to 4.5 nm. The latter is similar to that observed for the DDM sample and agrees with the I_f_ crystal prediction. We note that this crystal structure was obtained with a trapped ligand, deoxycholate. The recent work on MdfA in ND using NO spin labels reported no major changes for cytoplasmic pairs upon TPP binding. However, an increase in the maxima of the distance distribution of 0.6 nm was detected for the periplasmic pair TM3^101^ – TM11^373^ (12), which is on the order of the difference we observed for TM5^168^ – TM10^310^ in ND. To summarize this part, we detected structural differences in the periplasmic side, between ND and DDM environment in the absence of substrate, but not in the presence of substrate.

### Substrate binding

To determine the location of TPP in MdfA solubilized in DDM or reconstituted into ND, we employed an NO spin-labeled substrate, mito-TEMPO, which is an analog of TPP. (Fig. 1). The inhibition constant, *K*_i_, of mito-TEMPO/MdfA (wild type) complex in DDM was determined by competitive binding assay where the binding of [^3^H]TPP was inhibited by Mito-TEMPO (Fig. S4). Measurements were done with 1 μM of [^3^H]TPP in the presence of increasing concentrations of mito-TEMPO. The resulting *K*_i_ = 8.86 μM (RMSE=0.0047) is similar to the *K*_d_ of TPP (∼4.7 µM) (16). For this *K*_i_ and for the range of MdfA and mito-TEMPO concentrations (20-50 µM) used, we calculated that 52-66% of the MdfA proteins should carry a bound mito-TEMPO. We also verified the binding by continuous wave (CW) X-band (∼9.5 GHz) EPR. The addition of mito-TEMPO to Gd-C2 TM5^168^ MdfA resulted in the appearance of a slow motion component in the EPR spectrum which is indicative of binding (Fig. S5A). Simulation of the EPR spectrum of TM5^168^/mito-TEMPO 1:1 molar ratio in DDM showed that only 17-20% of the mito-TEMPO was bound (Fig. S5B). This suggests that the dissociation constant for the CL-MdfA Gd-C2 labeled mutant is higher than that of WT MdfA. Somewhat higher dissociation constants for TPP were already reported for MdfA mutants(2, 11).

We tested whether MdfA form oligomers by performing Gd(III)-Gd(III) DEER measurements on the singly labeled constructs in the absence of mito-TEMPO and observed only background decay with no modulation (Fig. S6A). To optimize the modulation depth, we tested samples with 1, 2, and 5 equivalents of mito-TEMPO (Fig. S6B) and the 1:2 ratio gave the largest modulation depth without introducing a too large background decay. This ratio was further used for all samples. Here we stress that carrying out Gd(III)-NO DEER under conditions of low binding is highly advantageous as compared to Gd(III)-Gd(III) or NO-NO DEER because of the spectroscopic selectivity feature the former offers. This allows increasing the relative substrate concentration to increase the % of bound MdfA, without paying the penalty of reduced DEER efficiency owing to a large background signal from the unbound subtract, which contribute to the observed echo and reduces significantly the modulation depth. In the case of Gd(III)-NO DEER, the NO signal does not contribute to the Gd(III) observed signal and its excess just increases the background decay.

For locating the position of the mito-TEMPO in the MdfA structure using the DEER derived distance distributions in DDM and ND we used triangulation, following the approach of Freed(22), where a bound lipid substrate was located in soybean seed lipoxygenase-1, and of Mchaourab and coworkers who located a spin-Labeled substrate in the multidrug antiporter NorM. (23) In these cases, both the substrates and the proteins were labeled with NO spin labels, while in our work the protein was labeled with Gd(III) and the substrate with NO, and Gd(III)-NO distance measurements(24-28) were carried out. We chose to label the periplasmic side, where we observed differences in the Gd(III)-Gd(III) distances between the two environments and where the binding of TPP led to a change in this distance (see Fig. 3). The Gd(III)-NO DEER form factors and the corresponding distance distributions of the three periplasmic singly labeled MdfA mutants TM3^101^, TM10^310^ and TM5^168^ with mito-TEMPO are shown in Fig. 4. In these experiments the observe pulses and repetition rates are optimized for Gd(III) (28) (see Fig. S2) to minimize NO contributions to the observed echo. For all samples, we observed a distance distribution with a major peak at a short distance in the range of 3-3.5 nm and a smaller peak at a longer distance of 4-5 nm. In general, the long distance peak is more intense in the ND samples. A small difference between the DDM and ND samples was detected only for TM5^168^, where the 3.5 nm peak in the DDM sample shifted to 3.8 nm in the ND sample.

**Figure 4.**
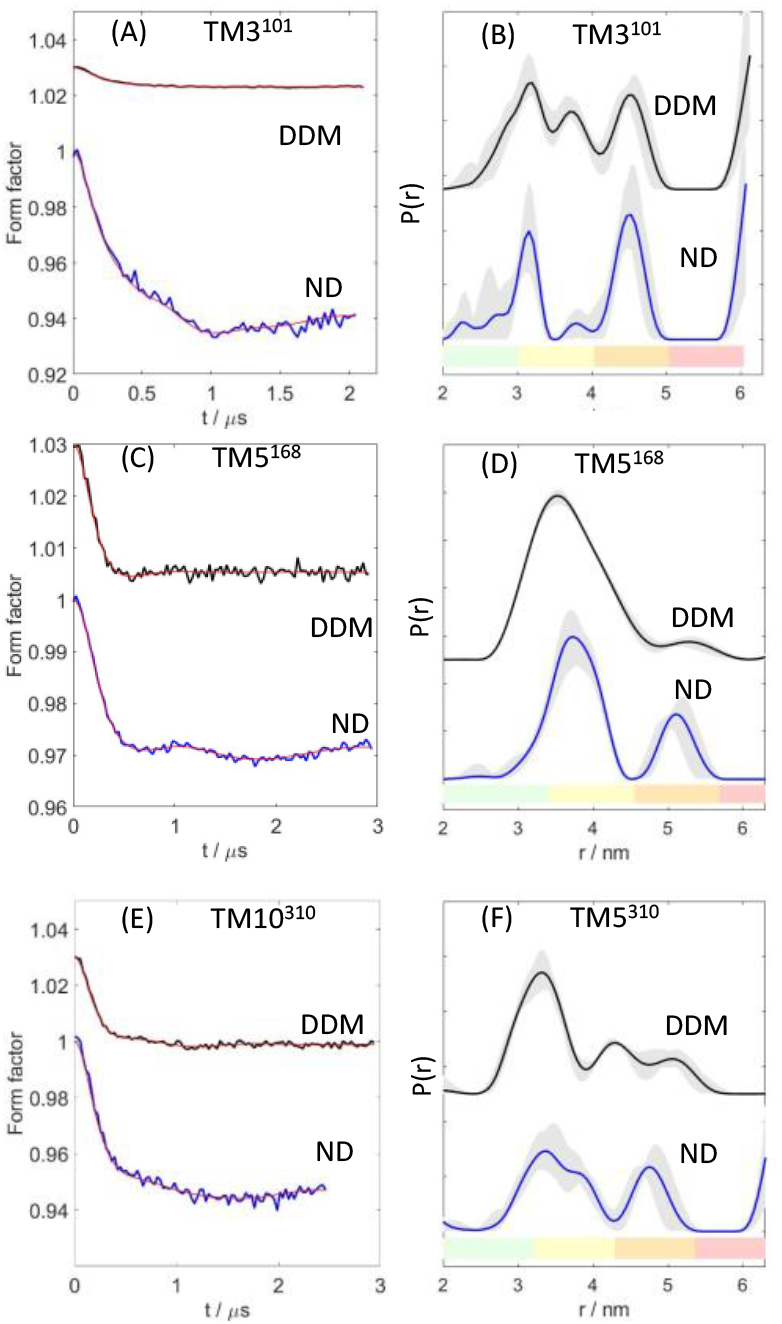
Gd(III)-NO DEER results of (A,B) TM5^168^, (C,D) – TM3^101^ and (E,F) TM10^310^ in DDM and ND with mito-TEMP. (A,C,E) The DEER form factor and (B,D,F) the corresponding distance distribution with color coding of the reliability regions as defined in DeerAnalysis. (Pale green: the shape of the distance distribution is reliable. Pale yellow: the mean distance and distribution width are reliable. Pale orange: the mean distance is reliable. Pale red: long-range distance contributions may be detectable, but cannot be quantified.) The light grey regions, representing the uncertainty range of the data, show alternative distributions obtained by varying the parameters of the background correction as calculated by the validation tool in the DeerAnalysis software package. The primary DEER data are shown in Fig.S7.

The modulation depth observed is only a few percent (2-6%), with the exception of the TM3^101^ DDM sample that showed a much lower modulation depth. This is significantly lower than the 12-15% expected for the experimental set up used, based on other samples measured in the lab. For the *K*_i_ determined above and a MdfA/mito-TEMPO molar ratio of 1:2 we expect that 74-87% of the MdfA proteins would carry a bound mito-TEMPO. This should yield a modulation depth close to 10%. If we consider the % of bound MdfA for 1:1 MdfA/mito-TEMPO derived from the EPR simulations, for a 1:2 ratio we expect only 31-37% bound MdfA, which is in better agreement with the low modulation depth observed. In general, we observed a higher modulation depth for the ND samples, which seems to correlate with a more intense long-distance peak. This maybe an indication for second binding site of mito-TEMPO within the lipid layers.

To validate that the DEER effect observed is indeed due to mito-TEMPO binding to MdfA, we added 10 equivalents of the stronger binder TPP to TM5^168^ /mito-TEMPO (1:2) and TM3^310^/mito-TEMPO (1:2). After 15 minutes of incubation we observed no DEER effect, except for the background decay (Fig. S6C), confirming that the distance distributions observed are indeed due to the interaction with the mito-TEMPO. Finally we exclude any contributions of NO-NO DEER to the distance distribution; although the observed pulse frequency overlapped with the NO signal (see Fig. S2), because of the saturation of the NO signal owing to the fast repetition rate and the pulse length optimized for the Gd(III) signal.

### The location of the mito-TEMPO in the MdfA substrate recognition pocket

From the Gd(III)-NO distance distributions for TM5^168^, TM3^101^ and TM10^310^, shown in Fig. 4, we determined the mean location of the NO group in mito-TEMPO in the I_f_ and O_o_ structures by triangulation(22), (23). Three distances is the minimal requirement for triangulations, naturally the larger the numbers of distances the higher is the accuracy.(29) Such calculations require the position of the Gd(III) within the protein structure and this was determined by modeling, where the Gd-C2 tag was grafted on the available crystal structures as described in the experimental section. First, we determined the mean position of the Gd(III) in each of the labeling sites, shown as cyan, green and orange spheres in Fig.5. Details, along with X, Y, Z coordinate’s distributions of the position of the Gd(III) ions for each tag conformation are given in Fig. S8 for the I_f_ structure and in Fig. S9 for the O_o_ structure. Using these distributions we determined the mean X, Y, Z coordinates for each Gd(III) labeling site and its mean location. These, and the Gd(III)-NO most probable distances (maxima of the distance distribution) were then used to determine the NO location by triangulation. For TM3^101^, where the intensity of the short and long distances are comparable, we took the peak of the short distance. Fig. 5 shows the two crystal structures, O_o_ and I_f_, with the mean positions of the three Gd(III) labels and the location of NO group of mito-TEMPO for ND and DDM. In the I_f_ structure, the most probable location of the NO group is close to the position of the deoxycholate ligand. The difference between the two environments is small, with the ND position situated on top of TM11, further away from the deoxycholate location. In Fig. 6A we show the location of the NO group with respect to the I_f_ structure with different ligands (deoxycholate and N,N-Dimethyldodecylamine N-oxide (LDAO)) and a substrate (chloramphenicol), where the proximity of the mito-TEMPO to the location of these compounds is obvious. In Fig. S10A we present a close up on the location of the NO in DDM and ND within the I_f_ structure highlighting the amino-acids in its close vicinity, that are mostly aromatic. In contrast, the location of the NO radical in the O_o_ structure is closer to the bottom of the open funnel, where it is surrounded by a smaller number of amino acid residues with which it can interact (see Fig. S10B). These results suggest that the I_f_ structure describes better the mito-TEMPO bound structure of MdfA than the O_o_ structure. The closer location of the NO radical to the substrates in the I_f_ structure for the DDM sample as compared to the ND samples indicates that the I_f_ structure reflects better the DDM solution structure than the ND structure.

**Figure 5.**
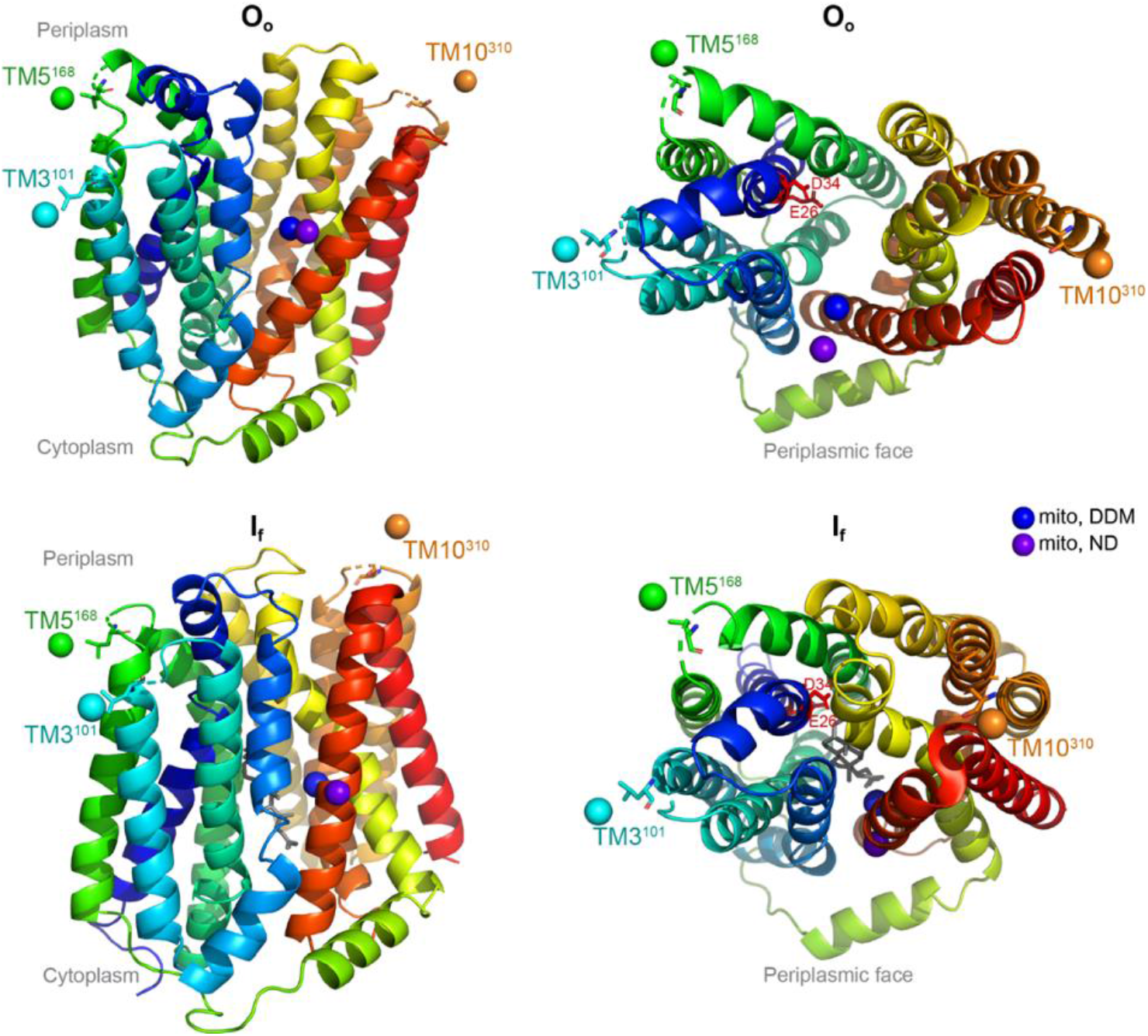
Two views of the location of the NO group of mito-TEMPO in ND (purple sphere) and DDM (blue sphere) in the O_o_ (top) and I_f_ (bottom) structures. The NO location was determined from the maximum of the short distance peak in the corresponding distance distribution. The views on the left include residues E26 and D34.

**Figure 6.**
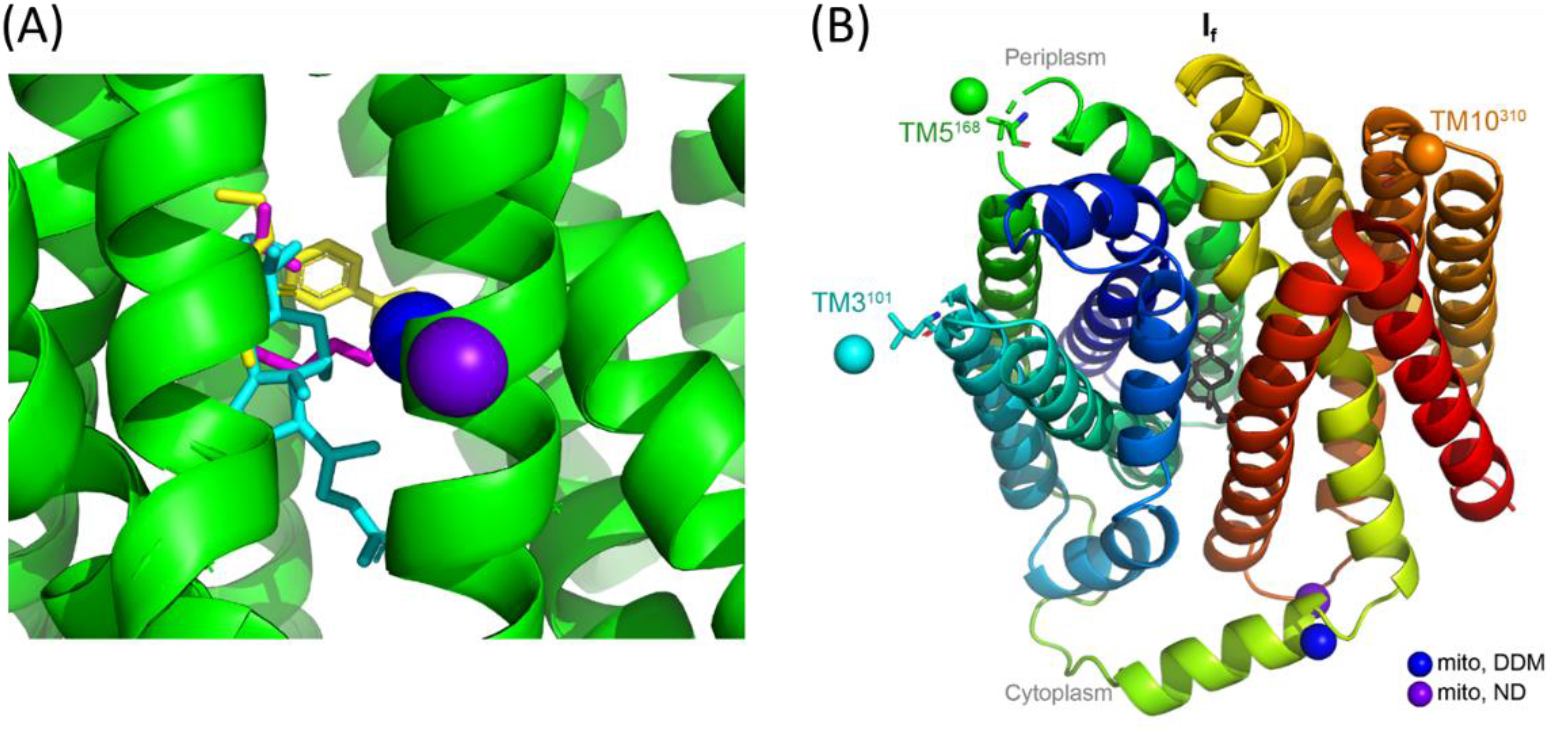
(A) Zoom on the aligned I_f_ MdfA structures with deoxycholate (PDB 4ZP0, cyan), dimethylamine N-oxide (PDB 4ZP2, magenta), chloramphenicol (PDB 4ZOW, yellow) and the location of the NO group of mito- TEMPO in ND (purple sphere) and in DDM (blue sphere). (B) The location of the NO group of mito-TEMPO in the second binding site in ND (purple sphere) and DDM (blue sphere) corresponding to the long distance in the distance distribution in the I_f_ structure.

We also evaluated the probability distribution of the NO location in the I_f_ structure, taking into account the width of the distance distribution peaks and the distribution in the Gd(III) location due to the flexibility of the Gd(III) tag (see SI for details). The result, shown in Fig. S11, reflects a rather large anisotropic distribution featuring a larger spread perpendicular to the membrane axis, which could be biased by the choice of the Gd(III) spin labels.

The Gd(III)-NO distances distributions of the ND samples showed an additional peak with a significant intensity at 4.5-5 nm, (Fig. 4), which suggests the presence of a second binding site for mito-TEMPO. Using these three distances, we located this second binding site of NO radical in the I_f_ structure at the entrance to the cytoplasmic side, just at the edge of the protein (see Fig. 6B). The Gd(III)-NO distance distributions of the DDM samples have peaks at the 4.5-5 nm range, which give an average location similar to that of the ND samples. However, the intensities of these peaks in the DDM samples are lower than in the ND samples, indicating a significantly lower population and higher uncertainty.

## Discussion

In this work we focused on substrate binding to MdfA solubilized in DDM, or reconstituted into ND, to obtain insight into the conformational changes it may undergo upon substrate binding and its sensitivity to its surrounding media, considering that ND are a superior membrane mimic environment than detergent micelles.(29-33) DEER studies, which compared these two environments using NO spin labels on various proteins, showed varying results on whether the ND reconstitution changes the conformation of the protein.(32, 33) It has also been shown that other parameters such as solvent or ligand accessibility might be enhanced in ND.(34) For substrate free MdfA in ND it was reported that the cytoplasmic side is closed, similar to that of the I_f_ structure, and somewhat less than in the O_o_ structure, which is in general similar to the substrate-free MdfA in DDM. (7, 12) Larger differences were observed for the periplasmic side, where in ND it is more closed than in DDM. For the former it was more similar to the I_f_ structure, whereas in DDM it resembled better the O_o_ structure, though somewhat more closed. (7, 12). Accordingly, the predominately outward facing O_o_ structure in DDM changes under the influence of the lipids to outward closed/inward closed (O_c_/I_c_) conformation.(12) The question that arises is whether the substrate location is also affected by the conformational changes induced by the environment on substrate free MdfA.

To determine the location of the substrate we made use of Gd(III)-NO measurements(24, 35, 36), between Gd(III) singly labeled MdfA and a NO labeled TPP analog, mito-TEMPO. The choice of employing Gd(III)-NO measurements, as opposed to the more standard NO-NO distance measurements allowed us to work under conditions of excess substrate with minimal interferences from the unbound substrate and avoid potential contributions of substrate-substrate distances to the distance distributions because the NO signal is saturated under the measurement conditions optimized for Gd(III) detection(28). In general, the Gd(III)-NO distance distributions observed for the three Gd(III) labeling sites were rather similar in DDM and ND (at neutral pH). The differences were limited to a somewhat shorter distance (0.3 nm) for TM5^168^ in DDM, larger modulation depth and the more intense peak at 4.5-5 nm for ND samples. For the DDM data, the average location of the NO radical of mito-TEMPO agrees rather well with the location of the ligands in the I_f_ structure (see Fig. 6a), suggesting the tri-phenylphosphonium part of the mito-TEMPO molecule is projected away from TM11 towards the substrate binding pocket. The position of the NO group, calculated from the ND distance distributions, gave an average position close to that of the DDM data (at a distance of 0.47 nm) but more pushed into TM11. This indicates that the I_f_ structure represents slightly better the DDM-solubilized MdfA with bound substrate than the ND-reconstituted transporter. The location of the NO group in the O_o_ structure seems to be more at the edge of the protein for both the DDM and ND samples and therefore we conclude that the O_o_ structure does not represent satisfactorily the occluded conformation of substrate-bound MdfA, in agreement with a recent study. (12)

We also calculated the distribution of the NO location in the structures, taking into account the distribution of the calculated Gd(III) positions in the structure arising from the flexibility of the Gd-C2 tether to the protein and the width of the distance distribution. The distribution was large and anisotropic, with the largest distribution found along the membrane axes. We attribute this mainly to experimental error in combination with choices of labeling sites, which were limited to the periplasmic side. The location of TPP within the I_f_ structure was also calculated using substrate docking.(8) In this work the I_f_ structure was used as the initial structure for MD simulations where the Q131R was abolished and docking of TPP was carried out on the resulting structures. The binding sites of TPP are distributed between the center of the recognition pocket, lined with and below residue D34, confined to the center of the MdfA, similar to the average position we obtained. The site distribution found in the docking calculations extended mainly along the channel perpendicular to the membrane axes towards the recognition pocket, which together with other residues, constitutes the rim around the entrance to the pocket from the cytoplasm. This rim comprises of E135, E136 R332 and E132 and the distribution indicated potential location around the rim, with rather low probability. This distribution was significantly narrower, than what we found.

Our results suggested the existence of a second binding site for the ND-reconstituted MdfA, as indicated by a second peak in the distance distribution at a longer distance. This site is located at the cytoplasmic side, near the edge of TM7. This position is however too far from the site in the rim indicated by the docking calculations. As these longer distance distributions were not prominent in the DDM sample, it is possible that the presence of the lipids in the NDs allowed the binding of TPP to this tentative second site. Recently, Yardeni et al.(12) identified a substrate-responsive lateral gate, in between TM5 and TM8 which is open toward the inner leaflet of the membrane and closes upon drug binding. Unfortunately, the site we have found cannot be assigned to this location as it is too far.

The Gd(III)-Gd(III) results, which track the MdfA conformational changes upon TPP binding, obtained under neutral pH and using different pairs than those described in ref (12), are consistent with the results of this work. For the cytoplasmic mutant, TM4^128^ – TM8^280^, the distance was not affected by the environment, whereas the periplasmic mutant TM5^168^ – TM10^310^ in apo-MdfA exhibited a significantly shorter distance in the ND, with a difference on the order of that observed earlier for TM3^101^-TM11^373^.(12) This earlier study showed that in ND, substrate induced conformational changes are small and limited to closure of the periplasmic edges of TM1 and TM7, increasing distance of the TM3-TM11 edges and shortening of TM5—TM10 and TM5-TM8 on the cytoplasmic side. These probably generate the occluded conformation, which does not involve rigid body movement of both halves (N6 and C6, representing the pseudo-symmetric halves of MdfA) against each other. Interestingly, upon TPP binding, the distance of TM5^168^ – TM10^310^ in ND increased, as observed for TM3^101^-TM11^373^, and it became similar to that observed in DDM, which did not changes upon TPP binding.

Several pairs involving labeling at the periplasmic edge of TM11 (TM11^373^ and TM11^371^) did not show any change upon TPP binding in ND (TM8^259^ -TM11^371^, TM5^163^ -TM11^373^, TM5^168^ -TM11^373^)(12). This indicates that change in distance is a consequence of the movement of TM3. Similarly, the change we observed for TM5^168^ – TM10^310^ and the invariance reported for TM5^163^ -TM11^371^ and TM5^163^ -TM11^373^ (12) suggests that the movement takes place at TM10. To account for the TM5^163^ -TM9^307^ small shortening (∼ 0.3 nm) with a concomitant increase of 1.1 nm for TM5^168^ – TM10^310^, upon TPP binding, TM10 should undergo a rotation. The TM3 and TM10 movement allows accommodating the drug and affords the closure of TM1 and TM7 to generate the occluded conformation (Fig. 8 in ref (12)). While there are clear differences between the conformations of apo-MdfA in ND and DDM, we did not detect differences in the substrate-bound occluded conformation. Interestingly, the position/orientation of TM10 in apo-MdfA DDM did not require any adjustment for the transition into the occluded conformation.

## Conclusions

Gd(III)-NO distance measurements between Gd(III) singly labeled MdfA and the TPP analog mito-TEMPO provided the location of the substrate within the I_f_ and O_o_ structure. The choice distance measurements between Gd(III) and NO, which display very different spectroscopic properties, allowed measuring DEER under conditions of a rather high dissociation constant, which require excess mito-TEMPO, with minimal interferences from the signal of the unbound mit-TEMPO. For both DDM and ND, the average position of the drug was found to be close to the ligand position in the I_f_ structure, with the DDM environment exhibiting a somewhat better agreement with the crystal structure than the ND environment. We therefore conclude that the I_f_ structure provides a good description of substrate bound-MdfA in DDM, but it is slightly modified in ND. A second substrate-binding site was found in the ND-reconstituted MdfA at the cytoplasmic side, towards the end of TM7.

In addition, Gd(III)-Gd(III) distance measurements on doubly labeled MdfA revealed conformational changes upon TPP binding. We detected significant differences in the periplasmic region in apo-MdfA, with ND featuring a more closed conformation than that observed in DDM. These findings are in agreement with recent reports where in ND apo-MdfA assumes an O_c_/I_C_ conformation, whereas in DDM the conformation resembles the O_o_ structure.(7, 12) The addition of TPP led to a noticeable conformational change in the periplasmic face in ND, attributed to a movement of TM10, increasing the TM5-TM10 distance. This is in addition to the movement of TM3 and closure of TM1-TM7 on the periplasmic side and an identified substrate lateral gate between TM5-TM8 in the inner membrane leaflet.(12)

The methodology presented here can be further applied to other MdfA drugs, having different chemical properties in order to resolve the question of how MdfA can accommodate chemically dissimilar substrates.

## Authors’ contributions

DG and EB designed research, TB and EHY performed research; DG and AF analyzed the data, DG, EB, EHY and AF wrote the manuscript. All authors read the manuscript.

## Acknowledgment

This research was supported by the Minerva Foundation (D.G.) and was made possible in part by the historic generosity of the Harold Perlman Family (D. G.). E.B. is supported by the United States - Israel Binational Science Foundation. We thank Prof. Thomas Huner, Australian National University for his help with the determination of the Gd(III) location within the available crystal structures. D. G. holds the Erich Klieger Professorial Chair in Chemical Physics. E.B holds the Ruth and Jerome A. Siegel and Freda and Edward M. Siegel professorial Chair

